# Comparative genomics of novel Bacteroides acidifaciens isolates reveals candidates for adaptation to host subspecies in house mice

**DOI:** 10.1101/2023.01.31.526425

**Authors:** Hanna Fokt, Shauni Doms, Malte C. Rühlemann, Maxime Godfroid, Ruth A. Schmitz, Britt M. Hermes, John F. Baines

## Abstract

The breadth of phenotypes influenced by the gut microbiome in multicellular hosts has attracted the keen and renewed interest of evolutionary biologists.

Comparative studies suggest that coevolutionary processes may occur as hosts and their associated microbes (i.e., holobionts) diverge. The majority of studies to date however lack information beyond that of 16S rRNA gene profiling, and thus fail to capture potential underlying genomic changes among microbes. In this study, we conducted a comparative genomic analysis of 19 newly sampled *Bacteroides acidifaciens* isolates derived from the eastern and western house mouse subspecies, *Mus musculus musculus* and *M. m. domesticus*. Through a panel of genome-wide association (GWAS) analyses applied to pangenomic content, structural gene rearrangements, and SNPs, we reveal several candidates for adaptation to the host subspecies environment. The proportion of significant loci in each respective category is small, indicating low levels of differentiation according host subspecies. However, consistent signal is observed for genes involved in processes such as carbohydrate acquisition/utilization (SusD/RagB, *amyA* and *amyS*) and de novo purine nucleotide biosynthesis (*purD*), which serve as promising candidates for future experimental investigation in the house mouse as a model of holobiont evolution.

## Introduction

The breadth of phenotypes influenced by the gut microbiome of mammalian hosts, and multicellular host-associated microbes in general (McFall-Ngai et al., 2013), has attracted the keen and renewed interest of evolutionary biologists. The term “holobiont,” which refers to a host organism and the entirety of its associated microbes (Gordon et al., 2013; Margulis and Fester, 1991), is recognized as an important unit of biological organization (Bordenstein and Theis, 2015). The conditions under which a host, microbe(s), or holobiont represent units of selection is an important area of research and a matter of discussion (Koskella and Bergelson, 2020; Moran and Sloan, 2015; Theis et al., 2016). However, as explained by Hurst (2017), the role of polymicrobial communities in organismal evolution may perhaps be best ultimately understood by considering “Darwin’s Tenets” at each level within a holobiont and identifying the forces that lead to variation in individual symbionts within and between hosts.

Comparative studies of the gut microbiome across hosts at various levels of evolutionary divergence have produced many intriguing observations. Phylosymbiosis, whereby the ecological relatedness of host-associated microbial communities parallels the phylogeny of their hosts, is observed in diverse host clades (Brooks et al., 2016). Alternative, but non-mutually exclusive processes could give rise to this community-level phenomenon, such as codiversification and/or coevolution versus related hosts being colonized by similar sets of microbes (Moran and Sloan, 2015). The observation of cospeciation across mammals for some individual bacterial taxa provides evidence for the former (Groussin et al., 2017; Moeller et al., 2016), for which coevolution may or may not be involved (Groussin et al., 2020; Moran and Sloan, 2015).

We previously employed the house mouse species complex to study the genetics of host-microbiome interactions between closely related host taxa, specifically the eastern and western host subspecies, *Mus musculus musculus* and *M. m. domesticus*, respectively (Doms et al., 2022; Wang et al., 2015). Interestingly, Wang et al. (2015) found that despite overall similar communities between the two subspecies, the genetic basis for maintaining gut microbial community structure likely diverged since their common ancestor ~0.5 million years ago (Geraldes et al., 2008). However, these—and related— studies were conducted using 16S rRNA/marker gene sequencing, which can obscure underlying bacterial genomic diversity (Douglas and Werren, 2016). Moreover, the 16S rRNA gene evolves too slowly to capture potential differentiation between closely-related host taxa. Thus, it remains unclear whether the divergence in the host genetic basis for gut microbial phenotypes is accompanied by differences in the pattern underlying bacterial genomic diversity, e.g., at the level of SNPs and/or their pan-genomes.

Here, we conducted a comparative genomic analysis of novel *Bacteroides acidifaciens* isolates derived from *M. m. musculus* and *M. m. domesticus* populations. *B. acidifaciens* commonly inhabits the mouse gut microbial community and is associated with diverse host phenotypes pertaining to health and physiology, including the prevention of metabolic disorders (Yang et al., 2017; Yang and Kweon, 2016), liver injury (Wang et al., 2022), and is also observed to display circadian oscillations (Thaiss et al., 2014). *Bacteroides* taxa display widespread host genetic signal for variation in their abundance (Rühlemann et al., 2021; Wang et al., 2015), including individual 16S rRNA gene amplicon sequence variants (ASVs) closely matching *B. acidifaciens* in our recent study of *M. m. musculus* and *M. m. domesticus* (Doms et al., 2022). Together with some signals of cospeciation at higher taxonomic ranks in other mammals [i.e. *Bacteroidaceae* (Moeller et al., 2016), Bacteroidales (Groussin et al., 2017)], we reasoned that *B. acidifaciens* represents an interesting model gut commensal to study bacterial genomic diversity according to host species, as comparable studies are largely limited to pathogenic taxa such as *Burkholderia* (Lee et al., 2021), *Campylobacter* (Costa et al., 2021), or *Staphylococcus* (Lima et al., 2022). Through a panel of genome-wide association analyses (GWAS) applied to pangenomic content, structural gene rearrangements, and SNPs, we reveal several candidates for adaptation to the host species environment, including genes involved in carbohydrate acquisition/utilization (SusD/RagB, *amyA* and *amyS*) and de novo purine nucleotide biosynthesis (*purD*), which may serve as a basis for future studies of coadaptation within the mouse holobiont.

## Results

### Sampling and genome sequencing of *Bacteroides acidifaciens* isolates

To obtain a set of representative *B. acidifaciens* genomes for comparative pangenome analysis, cecal contents were obtained from wild-derived outbred colonies of *M. m. musculus* [n=3 mice derived from Kazakhstan (KH)] and *M. m. domesticus* [n=3 mice derived from Massif-Central, France (MC) and n=1 from Iran (AH)] maintained in the same animal facility (Suppl. Figure S1). A total of 19 novel strains were isolated, ten from *M. m. musculus* from (KH), and nine strains from *M. m. domesticus* (MC, n=6; AH, n=3) (Supp. Table S1, Suppl. Fig. S1). Together with one representative reference genome, which was also isolated from mouse feces (RefSeq: MGBC000127; Figure 1A) (Beresford-Jones et al., 2022), we analyzed a total of 20 genomes. The *B. acidifaciens* genomes from our wild-derived mice show 99.84% pairwise average nucleotide identity (ANI) and 98.44% ANI against the reference genome. Genome sizes ranged from 4.62 Mb to 4.81 Mb (mean 4.68 Mb) and contained on average 4132 coding sequences (CDSs). In comparison, the reference genome is larger (5.2 Mb) and contains more CDSs (4596). A summary of the general characteristics of the genomes used in this study are provided in Supp. Table S1. Phylogenetic analysis of our *B. acidifaciens* isolates based on ANI in reference to type strains of five other *Bacteroides* species (NCBI) shows that they closely cluster with the *B. acidifaciens* type strain and are closely related to *B. caecimuris* (Fig. 1A).

**Figure 1.**
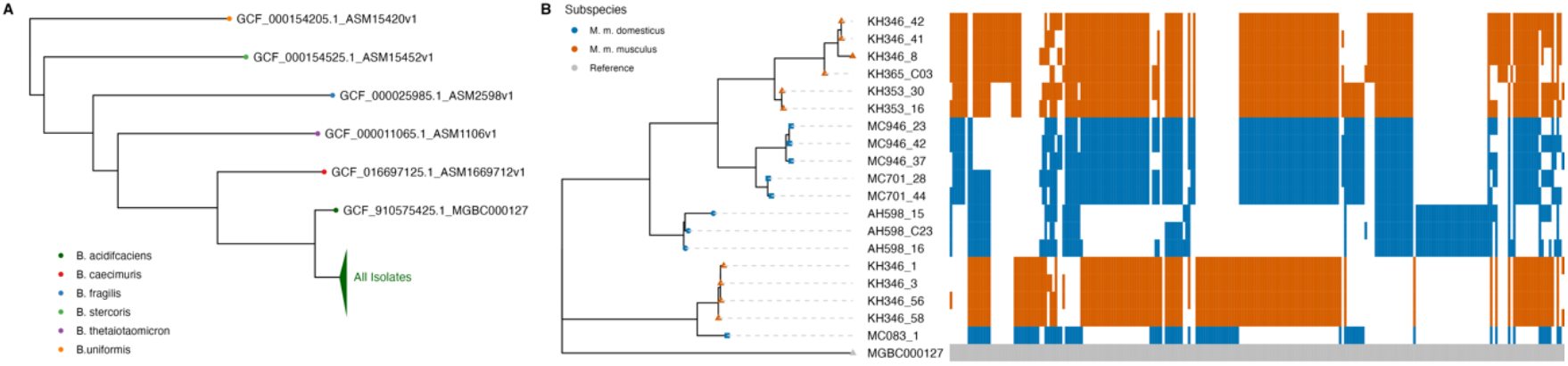
Phylogenetic overview of *B. acidifaciens*. (A) Global placement of *B. acidifaciens* reference sequence and isolates based on fastANI distances to five selected NCBI *Bacteroides* type strains. (B) Phylogeny of *B. acidifaciens* isolates based on core genome alignment. Presence/absence information of accessory genes is based on their location in the reference genome MGBC000127.

### Pangenome analysis

To infer the pangenome, we used Panaroo (Tonkin-Hill et al., 2020), which identifies 5,139 gene clusters including 2,750 strict core genes (present in all strains), 771 soft core genes (95% ≤ strains < 99%), of which 761 are present in all isolates but not the reference strain, 859 shell genes (15% ≤ strains < 95%), and 759 cloud genes (0% ≤ strains < 15%; Figure 1B; Suppl. Table S2). The soft core genes represent between 80.9% and 87.1% of a given genome. A maximum-likelihood (ML) tree based on the core gene alignment produced by MAFFT (Katoh, 2013; see Methods) reveals the *B. acidifaciens* isolates to display clustering according to individual mouse hosts (*λ*_pagel_=1, p=0.006), mouse origin (MC vs. AH vs. KH, *λ*_pagel_=1, p=0.0242) and a marginal effect of subspecies (*M. m. musculus* vs. *M. m. domesticus*, *λ*_pagel_=1, p=0.0612) (Fig. 1B). An exception is isolate MC083_1, a *domesticus-derived* strain that clusters with a set of divergent KH *musculus* isolates. From these genomes, we annotated a total of 5,139 genes using Prokka (Seemann, 2014).

We next annotated gene clusters with Cluster of Ortholog Genes (COG) categories using eggNOG-mapper (Cantalapiedra et al., 2021). The core gene clusters with known functions predominately belong to the categories ‘Carbohydrate metabolism and transport’ (G, n=278), ‘Cell wall/membrane/envelop biosynthesis’ (M, n=238), and ‘Replication and repair’ (L, n=235). Accessory gene clusters are principally classified as ‘Replication and repair’ (L, n=157), ‘Cell wall/membrane/envelop biosynthesis’ (M, n=103), and ‘Transcription’ (K, n=62). Eight COG categories are significantly differentially abundant between core and accessory gene clusters (Suppl. Table S3). The functional annotations of the predicted genes were not significantly enriched or depleted according to the originating mouse subspecies (Fisher’s exact test).

### Pangenome-wide association analysis

To determine whether variation in the *B. acidifaciens* accessory genome may be differentially associated with mouse subspecies, we performed a pangenome-wide association study (pan-GWAS) using pyseer (Lees et al., 2018). The analysis was based on 902 accessory gene clusters displaying presence/absence variation among our 19 strains. This reveals nine gene clusters to significantly associate with mouse subspecies (Bonferroni corrected *p* value threshold < 5.81E-4; Fig. 2; Suppl. Table S4), i.e., roughly 1% of the accessory genome. Interestingly, two sets of gene clusters have the same functional annotation, but are differentially abundant between the *domesticus*-and *musculus*-derived isolates. For example, gene cluster groups 932 and 939 are both classified as TonB-linked outer-membrane proteins in the SusC/RagA family; group 932 is predominately present in isolates from *M. m. domesticus* mice, whereas group 939 is present only in isolates from *M. m. musculus* mice. Similarly, groups 845 and 937 are both identified as belonging to the SusD/RagB gene family of outer membrane proteins involved in binding starch to the cell surface (KEGG orthology: K21572; Shipman et al., 2000). Group 845 is found in isolates mostly from *M. m. domesticus* mice; group 937 is present only in isolates from *M. m. musculus*. The remaining groups are *M. m. musculus*-associated, including group 826, which is annotated as a GTPase associated with the 50S ribosomal subunit, and group 1712 belonging to the ‘phage’ integrase family. Finally, group 1615 is annotated to COG category ‘S,’ a protein of unknown function, and groups 1790 and 1769 were not annotated.

**Figure 2.**
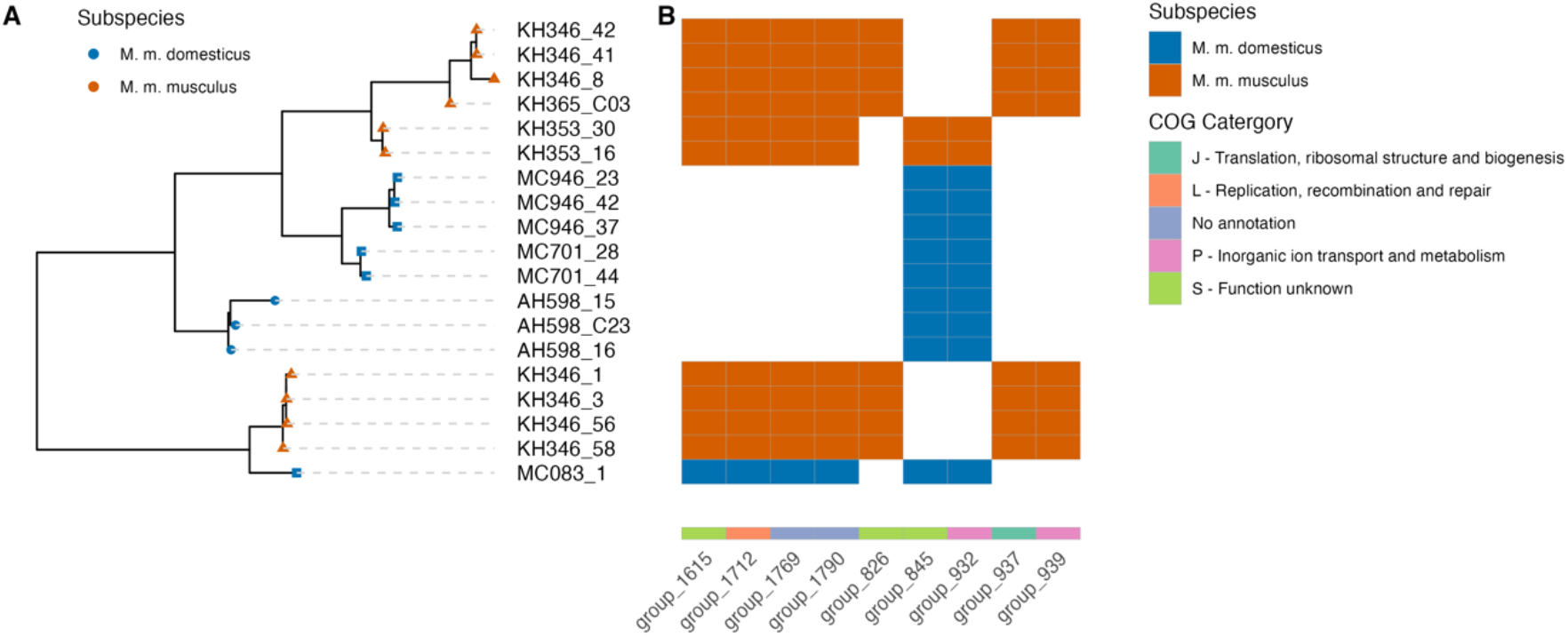
Significant gene clusters differentially abundant in strains isolated from different mouse subspecies using a pangenome-wide association study. (A) Phylogeny of *B. acidifaciens* isolates based on core genome alignment and presence in-formation of accessory genes. Labels on the y-axis represent the isolate ID and are colored to depict the mouse subspecies (blue: *M. m. domesticus*; orange: *M. m. musculus*); shapes represent the host’s geographical origin (triangle: Kazakhstan; square: France; circle: Iran). (B) Each block represents whether a gene cluster is present (colored) or absent (white). Gene cluster labels (x-axis) show the respective COG category.

### Structural variant GWAS (struct-GWAS)

As structural gene rearrangements are common in other *Bacteroides* (Carrow et al., 2020) and are known to associate with important phenotypes in such as antibiotic resistance (Tonkin-Hill et al., 2020), we next used Panaroo to detect structural variants. Rearrangements were analyzed at the level of consecutive gene triplets, and paralogs were split in the pangenome graph. Using the presence/absence of these triplets for genome-wide association analysis, we find ten significant structural variants out of a total of 296 that were detected (Bonferroni corrected *p* value threshold < 4.42E-4; Suppl. Table S5), which correspond to three genomic regions (Fig. 3). One large gene region with multiple structural variants includes a large insert before *mutS2*, a gene involved in the suppression of recombination (Pinto et al., 2005; Fig. 3A), and associates with isolates derived from *M. m. musculus* mice in addition to *M. m. domesticus* strain MC083_1 (Suppl. Table S5), which was previously shown to cluster with these same *musculus* strains (Fig. 1). We additionally find a region with two different structural variants around the genes *purD, ptpA*, and *rlmL* (Fig. 3B); this triplet significantly associates with *M. m. musculus* mice (Suppl. Table S5). Finally, we observe a region with one structural variant between the *yfnB* and *xerC24* genes (Fig. 3C; Suppl. Table S5).

**Figure 3.**
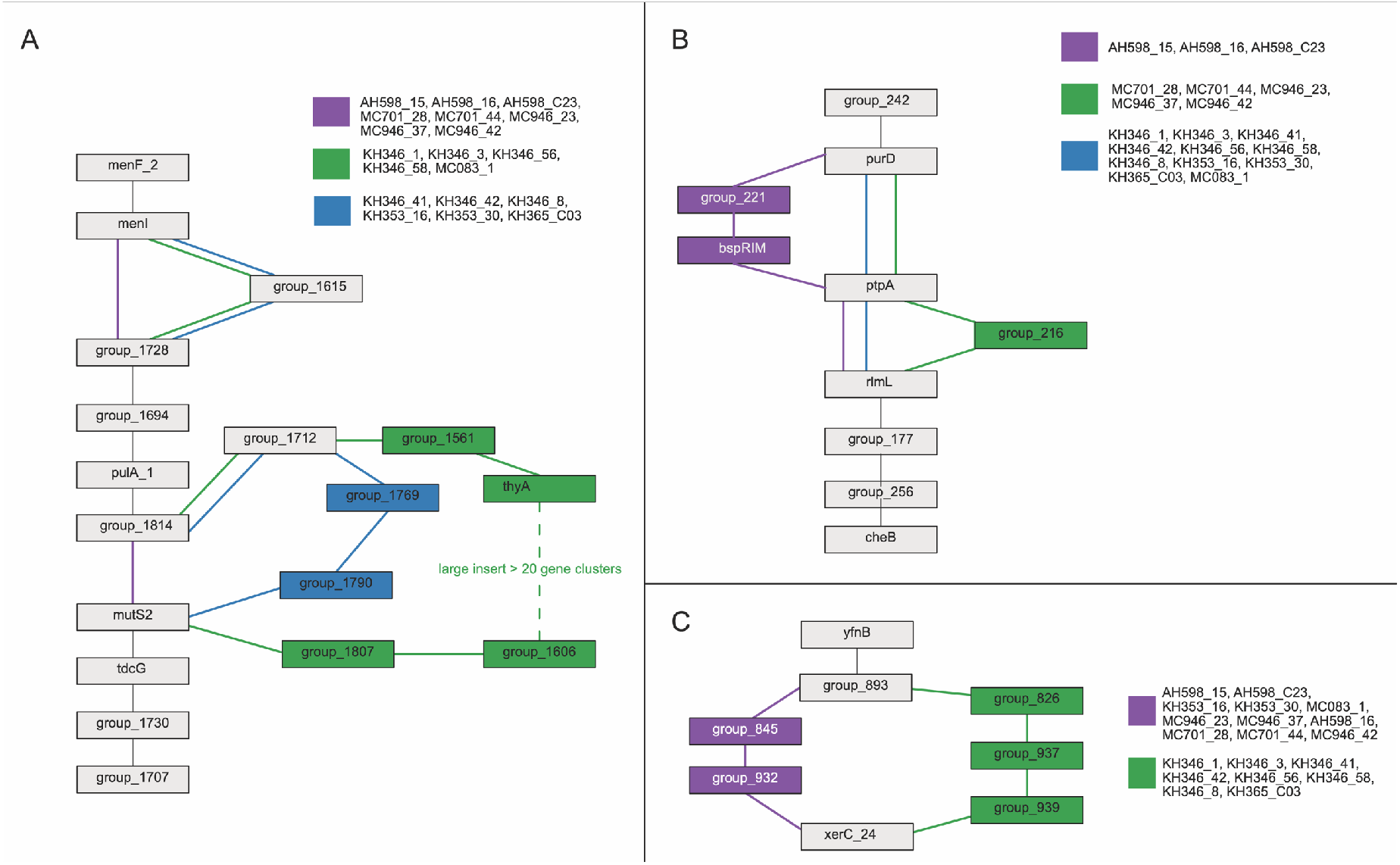
Structural variants associated with mouse subspecies. A complete representation of genes within the pangenome was constructed with Panaroo using gene rearrangements in the form of gene triplets (see Methods). The colors represent *B. acidifaciens* strains isolated from *M. m. musculus* or *M. m. domesticus*, originating from France (MC), Iran (AH), or Kazakhstan (KH), as specified in the isolate ID.

### SNP-GWAS

From the core genome alignment, we identified 15,168 single nucleotide polymorphisms (SNPs) among our 19 strains, which were included in GWAS analysis. We identify nine loci of interest (LOI) containing single nucleotide polymorphisms (SNPs) with signals surpassing the defined genome-wide significance threshold (Table 1; Fig 4; see Methods). Each locus spans about 200 kb from the first significant variant until the last significant variant (see Methods). SNPs within three loci (LOI 2, 3, and 9) contain no annotated genes. A complete list of genes included in these nine loci are provided in Supplementary Table S6.

**Fig. 4.**
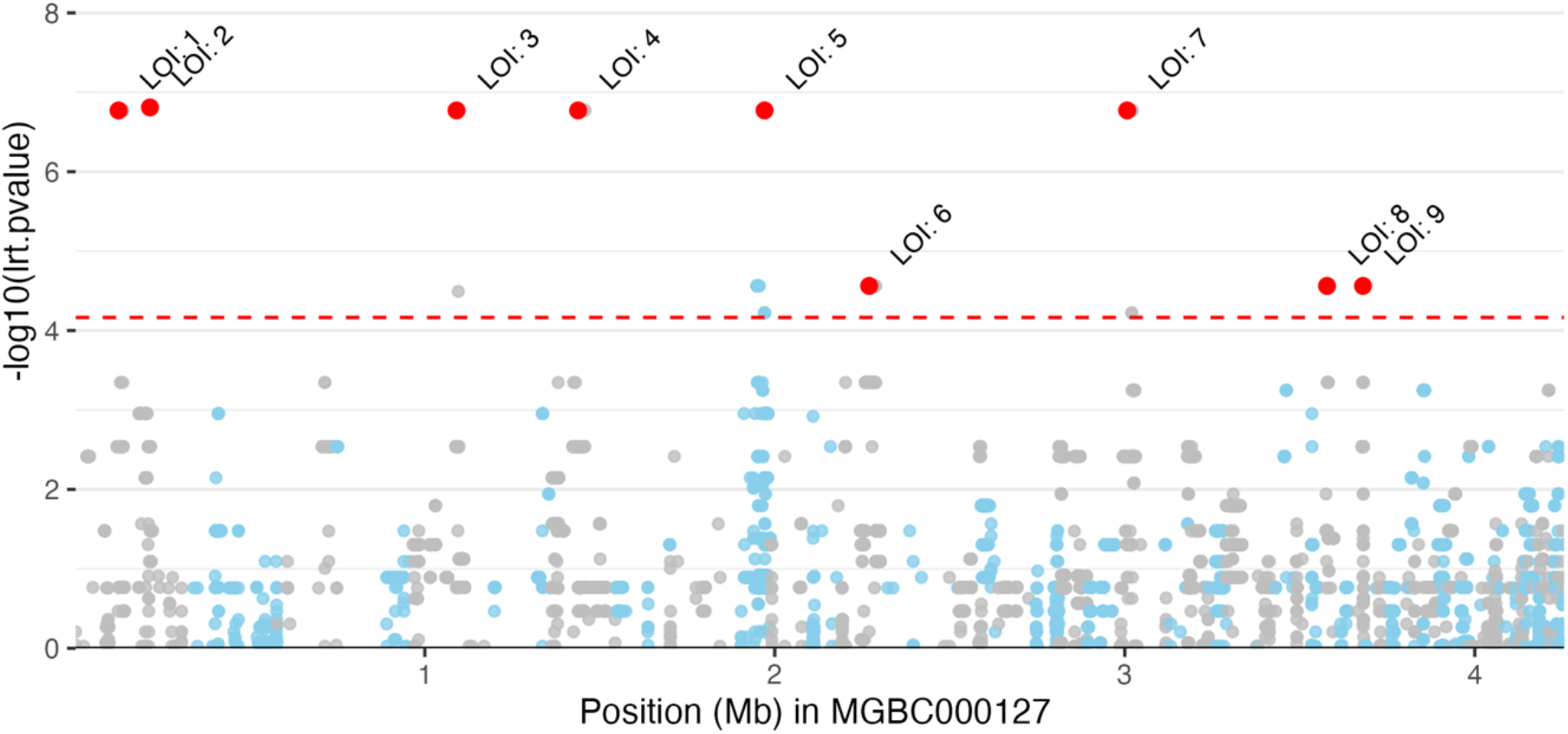
Manhattan plot of SNP-GWAS results; blue and grey colors alternate by contig, positions are continuous; lead variants per locus are marked. LOI = locus of interest. LOI numbers correspond with LOI presented in Table 1.

**Table 1.**
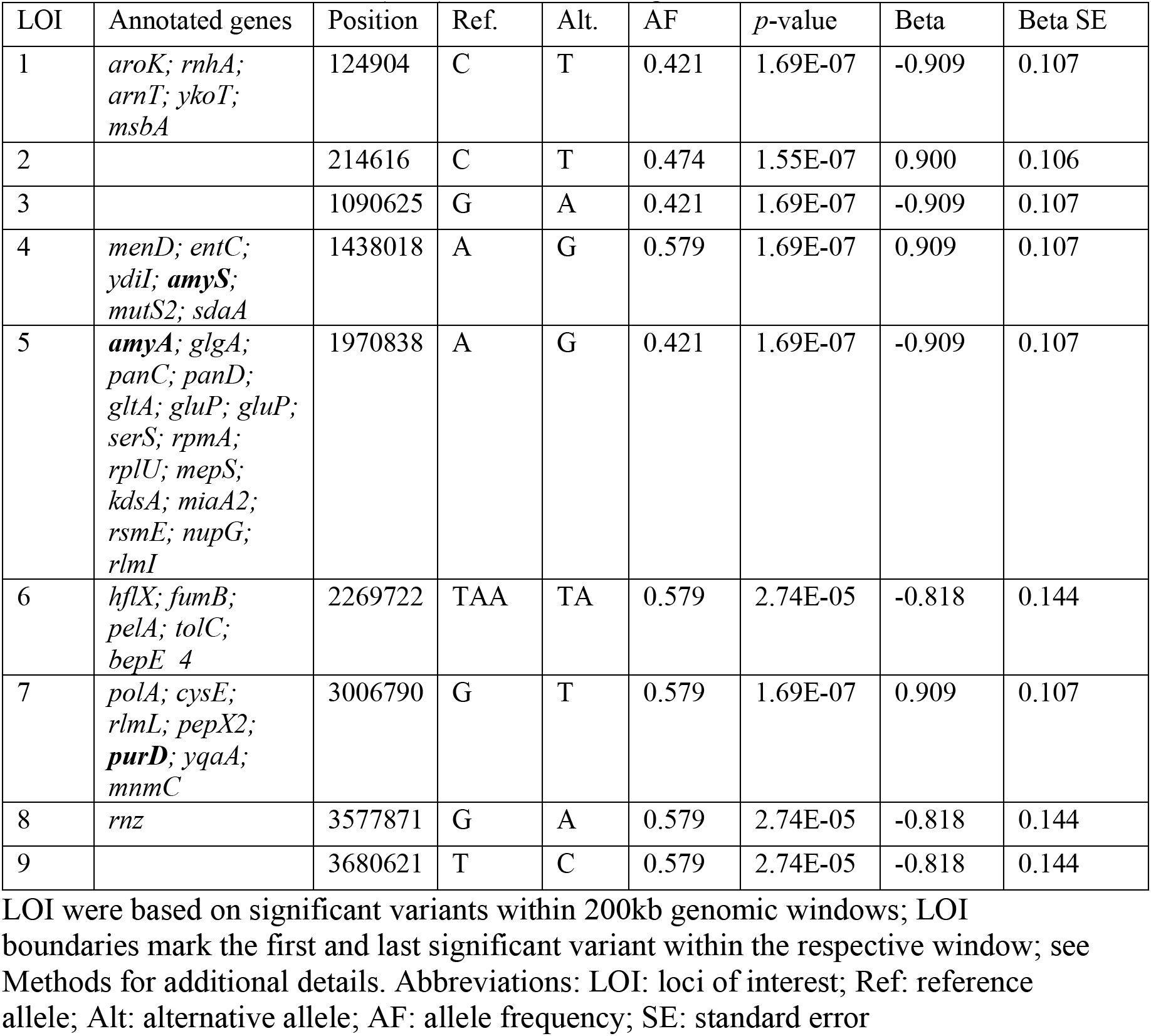
Nine loci of interest (LOI) with annotated genes.

Interestingly, we identified SNPs associated with several candidate genes previously detected in the struct-GWAS. For example, in LOI 1 we find *mutS2*, which was found to significantly associate with *M. m. musculus* (Suppl. Table S6. Similarly, we find *rlmL*, and *purD* in LOI 7, which were found to significantly associate with *M. m. domesticus* (Suppl. Table S6).

### F_ST_ analysis

Finally, to complement our panel of GWAS analyses, we calculated the fixation index (F_ST_) according to host subspecies in a gene-wise manner, as loci with F_ST_ values significantly higher than the genome average can indicate the action of population-specific selection. Candidate genes were identified as those displaying F_ST_ values >2.5 standard deviations above the genome average (Crits-Christoph et al., 2020). The average gene-wise F_ST_ among core genes is 0.216 ± 0.220 s.d., which indicates a substantial overall level of differentiation according to host subspecies. We find only one gene, *purD*, that exceeds the genome-wide threshold (*purD* F_ST_ = 0.793; cutoff = 0.767). Importantly, this gene was identified in struct-GWAS to be associated with the *musculus* subspecies and in the SNP-GWAS, within LOI 7 (Table 1; Suppl. Table S7).

## Discussion

Characterizing the nature of genomic variation in individual symbionts within and between hosts represents a key step in determining the role of polymicrobial communities in organismal evolution. Phenomena such as phylosymbiosis and cospeciation suggest that coevolutionary processes may occur between host and microbes, but in most cases information beyond that of 16S rRNA gene profiling is missing. In this study, we obtained whole genome sequence information from *B. acidifaciens*, a key gut taxon within the mouse holobiont. Through the analysis of roughly ten novel strains per mouse host subspecies, we identified a number of promising candidate genes that serve as examples of potential host-specific adaptation.

Interestingly, a number of the candidates we identified are involved in polysaccharide acquisition and metabolism. First, we observed that gene clusters including the SusC and SusD genes were differentially abundant between mouse subspecies. Starch utilization system (Sus) proteins form variable complexes to bind carbohydrates at the cell’s surface and utilize polysaccharides, and *Bacteroides* are highly dependent on the Sus system for nutrient acquisition (Martens et al., 2009; Tuson et al., 2018). Adaptive evolution involving the SusC/D genes in other *Bacteroides* taxa was observed even within the lifetime of individual human hosts (Zhao 2019), thus, it is not surprising that differences in these genes would be observed between host species. Additionally, our SNP-GWAS identifies the related amylase genes *amyA* and *amyS* as further loci of interest. Amylase is involved in the digestion of starches, and amylase copy number variation (CNV) is known to correlate with dietary starch intake in humans (Pajic et al., 2019; Perry et al., 2007). The mouse genome also displays substantial variation, including CNV and gain/loss events at the amylase gene cluster (Linnenbrink et al., 2020). It was previously hypothesized that human *AMY1* CNV may be involved in response to competition with commensal microbes over the utilization of starch (Walter and Ley, 2011). Thus, our observations suggest that a history of changes in diet and/or the host genome since the common ancestor of our sampled mouse populations may contribute to differentiation between *B. acidifaciens* strains.

Another interesting candidate revealed by our analyses is *purD*, which was identified in the struct-GWAS, the SNP-GWAS, and in the F_ST_ analysis. The *purD* gene is required for de novo purine nucleotide biosynthesis, which is conserved across the divisions of life (Yamamoto et al., 2022). Aside from being critical components of DNA and RNA, purine metabolites are involved in diverse signaling processes, in which the involvement of the gut microbiome is increasingly recognized (Li et al., 2022). Interestingly, humans carry a mutation in the *ADSL* gene that reduces the level of de novo synthesis of purines compared to other primates, suggesting that large-scale differences in purinergic signaling traits can evolve at the host level (Stepanova et al., 2021). De novo nucleotide biosynthesis is however also recognized as critical to the virulence of several pathogens, and may play a role in mediating competition between species (Goncheva et al., 2022; Truong et al., 2015; Yang et al., 2001). Thus, divergence in purinergic signaling between host and microbe, and/or differences in competition with other host subspecies-specific bacterial taxa, could contribute variation in *purD* among the *B. acidifaciens* strains in our study. Further studies exploring the role of *purD* in the growth and survival of *B. acidifaciens* in the mouse host are required to test this hypothesis.

While higher ranking bacterial taxa that include *Bacteroides* display some signatures of cospeciation, the signal is sometimes conflicting (Groussin et al., 2017; Moeller et al., 2016). For example, a more recent high-resolution analysis of human populations identifies both *Bacteroides* and the phylum *Bacteroidetes* to be among the taxa showing the least signal of codiversification (Suzuki et al., 2022). Thus, on the one hand, these results suggest that *Bacteroides* taxa may be rather viewed as generalist taxa among mammals, which would be consistent with the overall low number of significantly differentiated loci we observe according to host subspecies. On the other hand, our study is constrained by a relatively small number of bacterial genomes included in the GWAS analyses. Despite reduced power, we identified significant candidate genes that were consistent across several complementary analyses, suggesting a robust signal for these host subspecies-associated traits. The total number of significant loci identified in our study could thus represent a lower limit of the true number of host-associated bacterial genomic traits, whereby a wider sampling including additional genomes could reveal further candidates. Finally, it is also important to consider that the candidates observed here might be entirely related to the evolutionary history of *B. acidifaciens*, whereby other processes such as allopatric speciation (e.g., (Groussin et al., 2020)) could give rise to bacterial diversification between host subspecies.

In summary, our comparative analysis provides valuable insight into mammalian holobiont evolution in the house mouse model system, as it suggests that variation at the hierarchical level of the host can exert enough force to shape the pangenome even in a potentially widely dispersing, generalist species, and could thus still represent some of the initial stages of holobiont-level evolution. Future experimental and genetic interrogation of these candidate host-specific bacterial traits is thus clearly warranted.

## Materials & Methods

### Isolation of *B. acidifaciens* from cecal content

Cecal contents were obtained from the stock collection of wild-derived outbred colonies of *M. m. musculus* [n=3 mice derived from Kazakhstan (KH)] and *M. m. domesticus* [n=3 mice derived from Massif-Central, France (MC) and n=1 from Iran (AH)] maintained at the Max Planck Institute for Evolutionary Biology in Plön, Germany [for further details see (Harr et al., 2016)]. Mouse husbandry and sampling was conducted according to the German animal welfare law (Tierschutzgesetz) (Permit V 312-72241.123-34).

Fresh cecal content was placed in pre-reduced brain-heart infusion (BHI) containing 20% glycerol, homogenized, and stored at −80°C until further processing. Schaedler Anaerobe KV (SKV) Selective Agar with Lysed Horse Blood (Thermo Scientific) plates were purchased ready to use and stored at 4°C. Prior to inoculation, the SKV medium was reduced by placing the plates overnight under anaerobic conditions at room temperature. All the steps were performed under an anaerobic atmosphere (gas mixture: 5% H2, 5%CO2, 90% N2) in a vinyl anaerobic chamber (Coy Lab Products). Cecal contents were homogenized by vortexing and serial 10-fold dilutions in anaerobic BHI medium were made. Next, 50 μl of the last three dilutions (10^-6^, 10^-5^ and 10^-4^) were plated on SKV plates. The plates were incubated at 37°C for 48 hours. If the growth was insufficient after 48 hours of incubation, the plates were incubated for an additional 24 hours.

Colonies were picked from agar plates according to their morphology: *Bacteroides* form circular, white or beige, shiny, smooth colonies that are approximately 2-3 mm in diameter. Next, *Bacteroides* genus-specific primers Bac 32F and Bac 708R (Bernhard and Field, 2000) were used for the amplification of an approx. 750 bp portion of the 16S rRNA gene by colony PCR. PCR reactions contained 3.6 μl H_2_O, 5 μl of Multiplex PCR Master Mix (Qiagen), 0.2 μl of 2 μM Primer and 1 μl DNA template. Cycling conditions were as follows: initial denaturation for 15 min at 95 °C; 35 cycles of 30 sec at 94°C, 90 sec at 58°C, and 90 sec at 72°C, followed by 10 min at 72°C and ∞ 12°C.

Colonies confirmed to belong to *B. acidifaciens* by Sanger sequencing and matching to the RDP database training set 14 (Cole et al., 2014) were streaked on SKV agar plates. *Bacteroides* taxonomic status was double checked by PCR and Sanger sequencing as described above.

### Genome sequencing and processing

Genomic DNA for whole-genome sequencing was extracted using the DNeasy UltraClean Microbial Kit (Qiagen). Bacterial biomass from isolates grown on SKV agar plates was resuspended directly in 300 μl of PowerBead Solution (Qiagen) and vortexed to mix. All following steps were performed according to the manufacturer’s instructions. DNA samples were prepared according to Illumina Nextera XT protocol. The final DNA library was supplemented with 1% PhiX and sequenced on an Illumina NextSeq 500 system using NextSeq 500/550 High Output Kit v2.5 with 300 cycles.

Sequences were quality checked with FastQC (Andrews, 2010) and trimmed with Trimmomatic (Bolger et al., 2014) using a 4-base wide sliding window approach, cutting when the average quality per base dropped below 15. Reads with a minimum length of 40 bases were kept. Reads were assembled into contigs using SPADES (Bankevich, 2012). The quality of the assemblies was assessed with QUAST (Gurevich, 2013). Genomes were aligned to the RefSeq reference genome (MGBC000127, https://www.ncbi.nlm.nih.gov/assembly/GCF_910575425.1/) using bwa-mem (Li and Durbin, 2009) and further processed with samtools (Li, 2011) or variant calling. Filtering and quality control of the resulting file in variant call format (VCF) were performed with GATK (McKenna et al., 2010) and VCFtools (Danecek et al., 2011) applying the filters ““FQ < 0.025 && MQ > 50 && QUAL > 100 && DP > 15”.

### Genome annotation

Nineteen novel mouse isolates and the publicly available representative genome (MGBC000127, https://www.ncbi.nlm.nih.gov/assembly/GCF_910575425.1/) were analyzed similarly to reduce systematic errors. *De novo* gene predictions for coding regions were performed with PRODIGAL (v 2.6.3; (Hyatt et al., 2010)). Genomes were annotated using Prokka (Seemann, 2014). The pangenome calculation was performed using Panaroo (Tonkin-Hill et al., 2020). Additional functional annotation of the CDS was performed with eggNOG-mapper (v 2.1.6; (Cantalapiedra et al., 2021)) based on eggNOG orthology data (Huerta-Cepas et al., 2019). Sequence searches were performed using DIAMOND (Buchfink et al., 2015). Hypergeometric tests (Fisher’s exact test) were used to test for enrichment/depletion of functional annotations of predicted genes according to mouse subspecies. P-values were corrected for multiple testing using the FDR method.

### Phylogenomics

Core gene sequences of the isolated mouse strains were aligned using MAFFT (Nakamura et al., 2018); a maximum-likelihood tree was calculated using IQ-TREE (version 2.2.0; Minh et al., 2020; Chen et al., 2022) by applying the generalized time-reversible (GTR) model using 1000 bootstrap replicates for the ultrafast bootstrap feature (Minh et al., 2013). Pagel’s lambda (*λ*_pagel_; (Pagel, 1999) was calculated using the fitDiscrete() function in the geiger package for R (Pennell et al., 2014). P-values were calculated using a permutation approach using 1,000 randomizations of the traits (subspecies, origin, and individual) and comparison of the *λ*_pagel_ values from the randomized traits to the true traits.

### pan-GWAS, SNP-GWAS, and struct-GWAS

Pyseer (Lees et al., 2018) was used to perform GWAS analyses. The analysis was corrected for phylogenetic relatedness of the isolates by including a phylogenetic distance matrix calculated from the core genome alignment using the phylogenetic_distance.py script included in the Pyseer package. The number of unique patterns was tested using count_patterns.py, which uses this number to correct the *p* value using the Bonferroni correction method (Bonferroni, 1936). For the SNP-GWAS, loci of interest (LOI) were based on significant variants within 200kb genomic windows. LOI boundaries provided in Table 1 mark the first and last significant variant within the window. Supplementary table S6 provides the coordinates for all annotated genes in the reference genome.

### Calculation and analysis of fixation index (F_ST_)

F_ST_ for all variants were calculated in R (version 4.1; (R Core Team, 2022) based in the reference VCF file and using the vcfR package (Danecek et al., 2011). Using the gene boundaries in the reference genome, gene-wise F_ST_ values were calculated based on the ratio of the means of between- and within-population differences (Bhatia et al., 2013; Hudson et al., 1992). Only genes with at least 3 variant sites were kept in the analysis to identify genes exceeding the threshold of *mean*(*Fst*) + 2.5 × *s. d*. (*Fst*) = 0.24299 + 2.5 × 0.21661 = 0.78451.

## Acknowledgements

We thank Olga Eitel and Sven Künzel for technical assistance and Mathieu Groussin and Mathilde Poyet for comments on the manuscript. This work was funded by the Deutsche Forschungsgemeinschaft (DFG, German Research Foundation)—SFB 1182—project-ID 261376515 to JFB (Subproject A2) and RAS (Subproject Z2).

## Author contributions

J.F.B. conceived and designed the experiments; R.A.S. and J.F.B. supervised the research; H.F. performed the experiments; H.F., S.D., M.G., and M.C.R. analyzed the data; S.D., M.C.R., B.M.H., and J.F.B. prepared the original draft of the manuscript; all authors contributed to editing the final manuscript.

## Ethical approval

Mouse husbandry and sampling was conducted according to the German animal welfare law (Tierschutzgesetz) under permit V 312-72241.123-34.

## Data availability

This Whole Genome Shotgun project has been deposited at DDBJ/ENA/GenBank under the accession numbers JAQOUP000000000 - JAQOVG000000000. The version described in this paper is version XXXXXX010000000. The near full-length 16S rRNA gene sequences generated for this work have been deposited at the NCBI GenBank under accession numbers OQ312035-OQ312048. The draft genomes of the isolates have been deposited at NCBI under project number PRJNA926925.

## Code availability

Reproducible analyses and data tables used in this study are available at https://github.com/sdoms?tab=repositories.

