## Supplementary_figure_S1 for "Comparative genomics of novel Bacteroides acidifaciens isolates reveals candidates for adaptation to host subspecies in house mice"

### Supplementary Figures

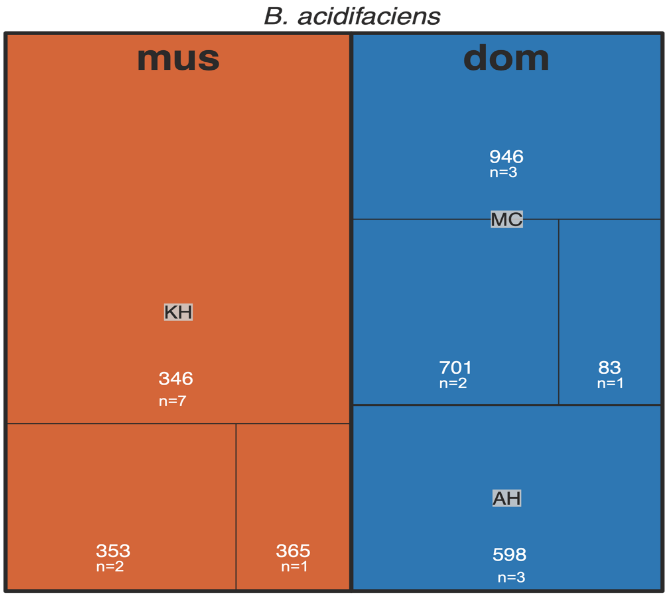


**Supplementary Figure S1**. Sampling overview of *B. acidifaciens* isolates. Ten isolates originate from *M. m. musculus mice* (n=3), originally captured in Kazakhstan (KH), nine isolates originate from *M. m. domesticus* mice, originally captured in Massif Central, France (MC, 6 strains isolated from 3 individuals), and Iran (AH, 3 strains isolated from one individual. Numbers indicate the mouse ID from where the strains were isolated.
